# Role of noise and parametric variation in the dynamics of gene regulatory circuits

**DOI:** 10.1101/291153

**Authors:** Vivek Kohar, Mingyang Lu

## Abstract

Stochasticity in gene expression impacts the dynamics and functions of gene regulatory circuits. Intrinsic noises, including those that are caused by low copy number of molecules and transcriptional bursting, are usually studied by stochastic analysis methods, such as Gillespie algorithm and Langevin simulation. However, the role of extrinsic factors, such as cell-to-cell variability and heterogeneity in microenvironment, is still elusive. To evaluate the effects of both intrinsic and extrinsic noises, we develop a new method, named sRACIPE, by integrating stochastic analysis with *ra*ndom *ci*rcuit *pe*rturbation (RACIPE) method. Unlike traditional methods, RACIPE generates and analyzes an ensemble of mathematical models with random kinetic parameters. Previously, we have shown that the gene expression from random models form robust and functionally related clusters. Under the framework of this randomization-based approach, here we develop two stochastic simulation schemes, aiming to reduce the computational cost without sacrificing the convergence of statistics. One scheme uses constant noise to capture the basins of attraction, and the other one uses simulated annealing to detect the stability of states. By testing the methods on several gene regulatory circuits, we found that high noise, but not large parameter variation, merges clusters together. Our approach quantifies the robustness of a gene circuit in the presence of noise and sheds light on a new mechanism of noise induced hybrid states. We have implemented sRACIPE into a freely available R package.

## Introduction

Noise or stochastic fluctuations in molecular components have been shown to play important role in many biological processes^1–3^, such as phenotypic switching, gene expression coordination, cell cycle and cell differentiation^4^, in both prokaryotic^5^ and eukaryotic organisms^3,4,6,7^. Noise can propagate in a gene network with a cascade of circuit motifs, and the expression level of a gene can vary up to six orders of magnitude between cells^1,8^. On the one hand, processes in gene regulation induce noise in the expression of transcripts or proteins, owing to factors such as transcription bursting and low copy numbers^1,8,9^; on the other hand, stochastic gene expression can influence the dynamics of biological systems^8^ or even create new dynamical features like oscillations, bistability etc^1,10–12^. It is not hard to imagine that, through evolution, cells eventually learn to use gene expression noise for their own advantage. For example, noise-induced cell-to-cell variability in protein levels in an isogenic cell population allows cells to assume different, functionally important and heritable fates^3^. This heterogeneity in clonal populations of cells may be essential for many biological processes as it enables the cells to respond differently to inducing stimulus^1^. Conversely, heterogeneous individuals in different environments can produce the same cell phenotype through phenotypic buffering/capacitance^8^.

Previous studies have unveiled many features of stochasticity in gene expression, yet, a comprehensive and quantitative understanding of the noise-induced dynamics is still elusive^13^. Various mathematical frameworks^14,15^ have been developed to model the dynamics of gene regulatory circuits (GRCs) governing cellular processes. Here, a GRC is a *functional* regulatory network motif, composed of a small set of interconnected regulators. To study the dynamics of gene circuits, various simulation schemes have been developed, including stochastic simulation algorithms (SSA, such as Gillespie algorithm^16^), methods solving stochastic differential equations (SDEs), asynchronous random Boolean network model^17^, hybrid methods that incorporate stochasticity in both discrete and continuous variables^18^, and the multiscale nature of different types of noise^19^. However, most of these methods require a fixed set of kinetic parameters that are associated with the regulation of individual genes, such as production rates, degradation rates, and binding/unbinding rates of protein-DNA (dis)association. Unfortunately, it is very hard to measure these parameters directly from experiments^14^, and therefore it limits the accuracy and prediction power of the traditional simulation schemes.

We have recently developed a systems-biology modeling method, named *<u>ra</u>*ndom *<u>ci</u>*rcuit *<u>pe</u>*rturbation (RACIPE)^20^, to deal with this long-lasting issue of parameter uncertainty. RACIPE takes the GRC topology as the only input, and generates an ensemble of models with random kinetic parameters. Then conventional ordinary differential equation based simulation is used for each random model to obtain steady-state gene expression. Finally, statistical analysis is performed on the *in silico* gene expression data from all the models to obtain the robust features. From our previous tests on simple GRC motifs and the biological regulatory circuits governing epithelial-to-mesenchymal transition (EMT)^20^ and B-cell development^21^, *etc*., we found that the steady-state solutions from an ensemble of random models form several distinct clusters according to their expression patterns, which correspond to the functional states of the circuits (e.g. the functional states A^ON^B^OFF^ and A^OFF^B^ON^ for a toggle switch with two genes A and B). The spread of the parameters of the models in a particular cluster can be associated with the robustness of the functional state against the parametric perturbation.

To facilitate the stochastic analysis, here we present a new modeling method that integrates stochastic methods with RACIPE. Compared to existing methods, this method has the following advantages. *First*, the stochastic RACIPE (sRACIPE) provides a holistic picture to evaluate the effects of both the stochasticity in cellular processes and the parametric variations. Typically the noise in cellular processes is regarded as “intrinsic” if it is caused by the stochastic nature of transcriptional, translational and post-translational regulations due to either low copy number of molecules or slow switching among the states of promoter structure, chromatin epigenetics, or nuclear architecture^8^. If the noise is due to pathway specific or global differences in the abundance of cellular components, or due to differences in the timing of cell-cycle events, it could be considered as “extrinsic”^22,23^. Segregating the effects of “intrinsic” and “extrinsic” noises in gene expression is not straight forward and is being actively studied^1,24^. Our randomization-based method, sRACIPE, captures the effects of both the intrinsic and extrinsic noises as it incorporates both the stochastic fluctuations and the parametric variations. *Second*, sRACIPE allows us to evaluate the effects of noise on the cellular states of a GRC. In conventional mathematical modeling, a cellular state is defined as a stable steady state (fixed point) of a nonlinear dynamical model. However, when the signaling of the system alters, the corresponding fixed point shifts accordingly. Therefore, it is particularly difficult to associate different steady states to a cellular phenotype. To deal with this issue, we define a distinct cellular state as one of the clusters of steady state gene expression profiles from random models^21^. With sRACIPE, we can evaluate how gene expression noise affects the formation of the clusters and the changes in their expression patterns. *Third*, the stochastic analysis can quantify the relative stability of the steady states for systems allowing multiple states. This is especially hard in the original RACIPE, where the deterministic analysis is adopted to solve the rate equations and where every stable steady state was considered equally probable.

To integrate the stochastic analysis with RACIPE, we have to address an important challenge as described below. Typically, one starts from an initial condition and runs stochastic simulation for a long time to obtain the steady-state probability distribution and transition rates. In RACIPE, we generate a large (∼10^4^-10^6^) number of random models and this stochastic simulation scheme will have a very high computational cost. Moreover, each model has a distinct set of kinetic parameters; therefore, the convergence of one model does not necessarily imply the convergence of another. A good simulation scheme has to be designed to reduce the computational cost without sacrificing the convergence of statistics.

In the following, we will introduce the stochastic analysis methods employed in sRACIPE. We will first describe two simulation schemes – a constant noise based method to estimate the basin of attraction of various states and another simulated annealing based method to compare the relative stability of different states and find the most stable state of GRCs. We will illustrate the methods using the canonical double-well potential system. Afterwards, we will explain the integration of these stochastic analysis methods with RACIPE and apply the new method on simple toggle switches, coupled toggle switches and an epithelial-mesenchymal transition (EMT) network^20,25^. We will demonstrate how the parametric variations and noise influence the functions of GRCs. The workflow of the sRACIPE method is presented in Fig.1.

**Fig. 1:**
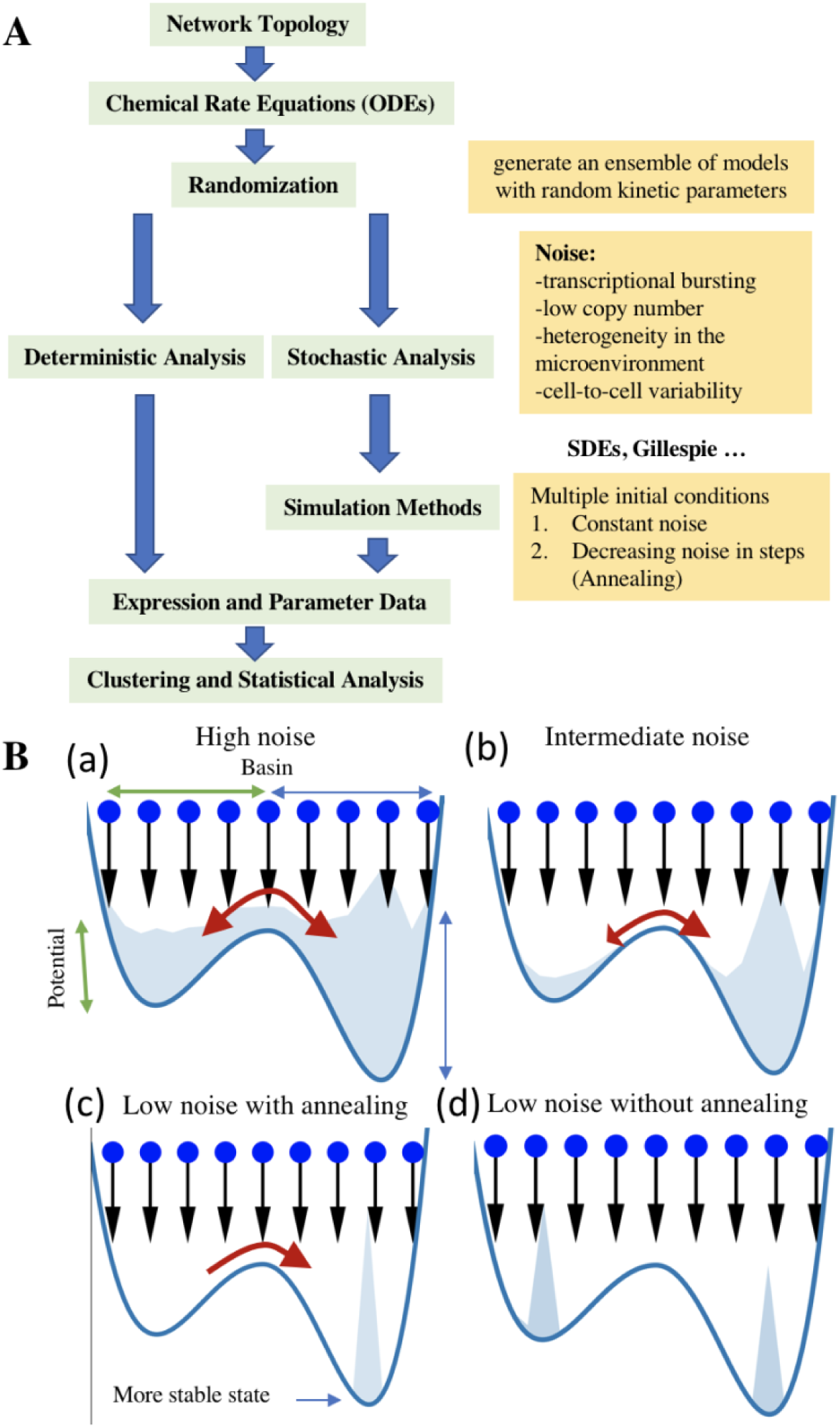
Illustration of sRACIPE. **(A)** The workflow of sRACIPE. The method integrates two ensemble-based sampling schemes – constant noise with multiple initial conditions (MIC) and simulated annealing (SA). **(B)** Illustration of the stability and basin of attraction using an example of a double well potential. (a) High noise enables frequent transitions between the minima, and the steady-state probability distribution in each well is proportional to the stability of the well. (b) Intermediate noise permits a larger number of transitions from the less stable well to the more stable well, and some trajectories are trapped in the more stable well. (c) As the noise is decreased further, the annealing based sampling scheme results in more occurrences of the particles in the more stable well. (d) For low noise cases, the transitions between wells are rare. Thus, each well traps all the particles in its basin of attraction, and the steady state probability distribution is proportional to the basin width of the wells.

## Results

### Sampling schemes for stochastic analysis

The temporal dynamics of a dynamical system can be obtained through numerical simulations of stochastic differential equations (SDEs) or Gillespie/kinetic Monte Carlo algorithms. A standard approach is to start with a random initial condition, run the simulation at a constant noise level for a long time, and record the state variables at equidistant time points. The histogram of these state variables gives the steady state probability distribution of the system. Here, we refer to this method as single initial condition (SIC) method.

For a system with multiple minima and a low noise level, the SIC method converges slowly as the system gets trapped in a local minimum^26^. To address it, we can instead perform statistics on an ensemble of simulations. Here, the method performs multiple simulations for a short simulation time starting from different initial conditions, and then it records the state variables only once at the end of each simulation. This approach, referred to as multiple initial conditions (MIC) method, has three advantages: (1) it can simultaneously sample multiple configurations of the system, therefore providing better coverage; (2) it can be naturally integrated into RACIPE as RACIPE is also an ensemble based method; (3) it can be easily parallelized as each initial condition evolves independently of others. Indeed, MIC and its variants^27,28^ have been adopted in simulations of equilibrium systems, but it is nontrivial for non-equilibrium systems^27^. However, in the low noise scenario, while MIC can sample multiple configurations (thus basins of attraction^29,30^), each of the trajectories is still trapped in a local minimum; therefore, it does not estimate the stability of the minima.

Here, we propose another sampling scheme based on simulated annealing^31^ (SA) to investigate the stability of a system. This method also generates an ensemble of simulations using multiple initial conditions. Each simulation starts with a random initial condition and a large noise level. Then, a constant noise simulation is performed for relaxation, and the state variables are recorded. The corresponding histogram of state variables from the ensemble of simulations gives the steady state probability distribution at that noise level. After the initial stage, the noise is reduced to a slightly lower level. Here, the states obtained from the simulations of the previous noise level provide a good estimate of the initial conditions for the simulations at the next noise level. This procedure is repeated till the system reaches zero noise. The simulations from the whole protocol produce steady state probability distributions at various noise levels. The initial high noise allows the simulations to adequately sample multiple minima, while the intermediate to low noise levels allow more transitions from less stable minima to more stable minima, eventually reaching the most stable state (Fig.1B). In the following sections, we will show how we tested the sampling schemes and how we integrated them into RACIPE to study the stochastic dynamics of GRCs.

### Comparison of the sampling schemes in double-well potentials

We first tested the three sampling schemes for stochastic analysis, *i*.*e*. SIC, MIC and SA, in the canonical double well potentials, where the corresponding potential functions can be obtained analytically. Calculation of such potentials for GRCs is usually difficult and computationally intensive^32,33^. Tests were performed on four variants of double-well potentials, where each variant differs from others in terms of the basin width and/or stability of wells. In SIC, the histogram of the particle positions was obtained from the positions at equidistant time points from a long simulation at a specific noise level. In MIC, the histogram was generated from the final positions of multiple short simulations for a fixed noise. In the SA scheme, histograms for different noise levels were obtained from the final positions of all the short simulations for the corresponding constant noises during simulated annealing.

For each potential variant shown in the first row of Fig. 2, the 2^nd^-4^th^ rows in the same figure show the corresponding steady state probability distributions at different noise levels using SIC (2^nd^ row), MIC (3^rd^ row), and SA (4^th^ row). At high (blue curves) and intermediate (orange curves) noise levels, the probability distributions from all the methods converge in all the four variants, as noise is large enough to induce sufficient transitions between the two basins. However, at low noise levels (green curves), a single trajectory is trapped in one of the basins. Thus, SIC, unlike the ensemble-based methods MIC and SA, never yields a converged distribution for all the four variants. For the fully symmetric double well potential (Fig. 2A, Fig. S1), both MIC and SA yield same probability distribution in all the cases. When the two wells have the same basins of attraction but different stability (potential), MIC provides equal probability in both wells but SA identifies the more stable well (Fig. 2B, Fig. S1). If the two wells differ in their basins of attraction but have same minimum potential values (Fig 2C, Fig. S1), the probability distributions obtained from MIC are proportional to their basins of attraction. However, SA has all the probability in the well with the larger basin. Lastly, when one well has larger basin width and the other is more stable (Fig. 2D, Fig. S1), SA correctly yields all the probability in the more stable well (supplementary video) whereas the probability distribution from MIC is proportional to the basin width. Altogether, our tests demonstrate that MIC and SA complement each other, especially for low noise cases, when MIC better estimates the basin of attraction and SA better estimates the stability.

**Fig. 2:**
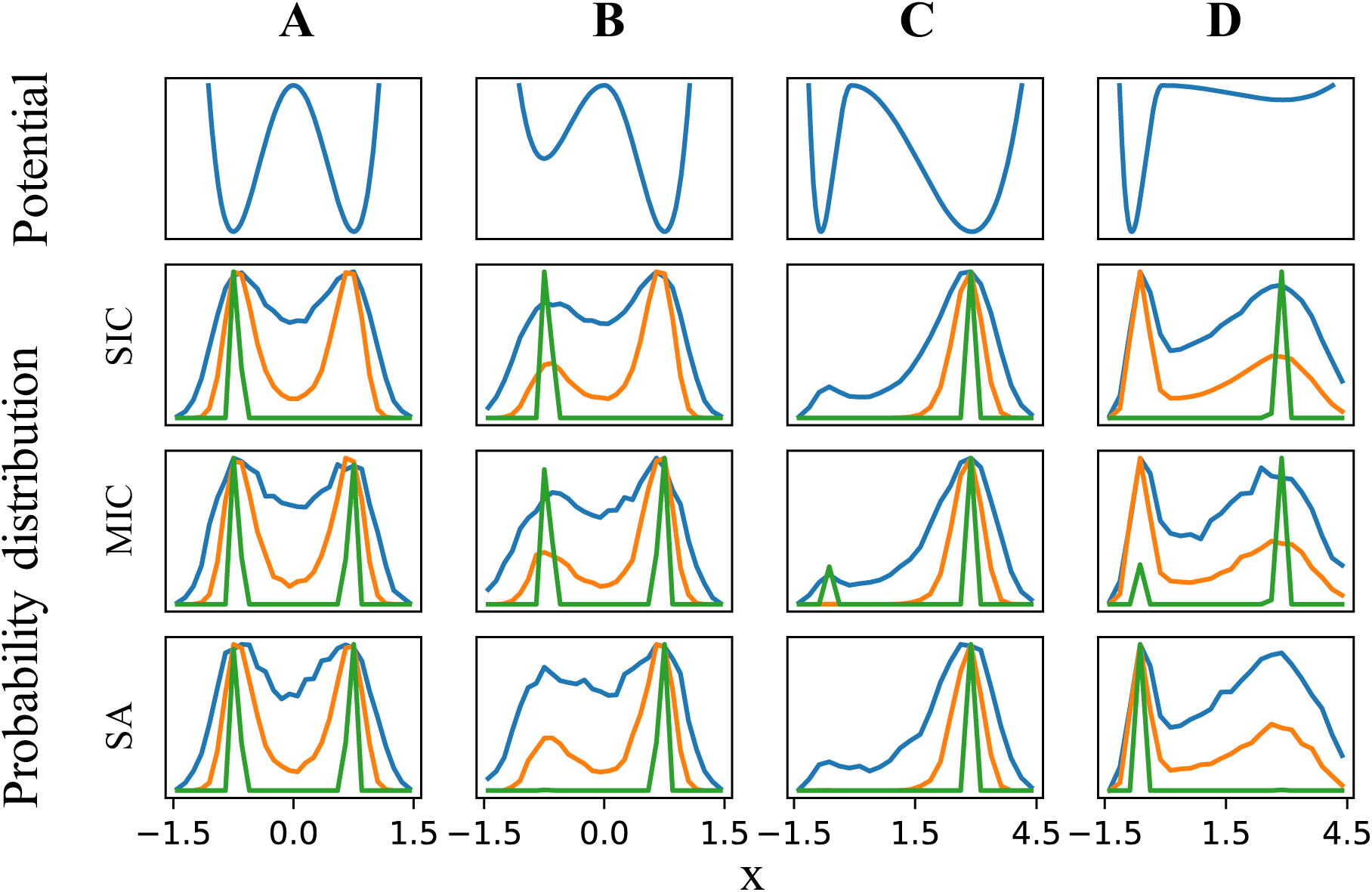
Tests of the stochastic analysis methods using double well potentials. First row shows four variants of double well potentials. (A) The two wells have identical potentials as well as the basins of attraction; (B) the left well has lower stability than the right well, but their basins are same; (C) two wells have same stability but asymmetric basins with the right well having a larger basin, and (D) the two wells differ in stability as well as basins such that the left well is more stable but has a smaller basin of attraction. The 2^nd^-4^th^ rows show the histogram of the steady states for different sampling schemes, namely, single initial condition (SIC, 2^nd^ row), multiple initial conditions (MIC, 3^rd^ row) and simulated annealing (SA, 4^th^ row) for each of the four potentials and at three different noise levels - high (blue), intermediate (orange) and low noise (green). All of the methods converge for the high and intermediate noise levels whereas, for the low noise, the particle is trapped in a random well in the SIC scheme, MIC captures the basin of attraction of the wells and SA captures the most stable well. The equations, parameter values and figures for other noise levels (Fig. S1) are available in SI.

### Integration of stochastic analysis into RACIPE

In the above sections, we have described two ensemble-based sampling schemes for stochastic analysis. Here, we will introduce a new method named sRACIPE, which integrates these sampling schemes with RACIPE. In the case of double-well potentials, the simulations using multiple initial conditions in MIC and SA can be considered as simulations of an ensemble of identical models using only one initial condition for each model. In contrast, the models in sRACIPE are not identical as it generates a large ensemble of random models, and each of these models is subject to a simulation scheme (either MIC or SA) using one initial condition only. We chose this scheme because of the following reasons. First, since sRACIPE generates a very large number of models, there are multiple models with similar parameters, and a collection of these models will identify most of the states. Second, as we learned from our previous studies, increasing the number of models provides better convergence of the probability distribution of the simulated gene expression data compared to increasing the number of initial conditions^21^. Third, we have tested and found similar results when sampling multiple initial conditions for each random model (Fig S4).

In the first MIC-based sampling scheme, for each model, a short simulation using a random initial condition and a fixed noise level is used to obtain the gene expressions of that model at that noise level. Such gene expressions from all the models are used for further statistical analysis. This procedure is repeated for other noise levels to obtain gene expressions for those noise levels.

In the second SA-based simulation scheme, we first pick a random initial condition for a model and perform a simulation at a high noise level. Then, for each model, using the final gene expressions from the simulation at a higher noise as the new initial condition, we perform another simulation at a slightly lower noise level. We repeat this procedure until the noise level gradually decreases to zero (details in SI). The final gene expressions from all the simulations and all the models are used for further statistical analysis for the corresponding noise levels. In the following, we will show the application of sRACIPE to several biological GRCs.

### Expression noise induces state merging

We applied sRACIPE to a toggle switch GRC consisting of two mutually inhibiting genes (Fig.3A, the rate equations shown in SI). Here, the MIC-based method was used to obtain the gene expression profiles for an ensemble of models. To obtain features at different noise levels, we considered the noise level as an additional model parameter and randomized it from a uniform distribution ranging from 0 to 50. Fig 3A shows the 2D histogram of the normalized gene expression at different noise levels. At low noise levels, we observe two distinct clusters or states, as evident from the histogram on the left showing the distribution of the expressions of gene A for noise levels between 0 and 1. The distribution is similar to that from the deterministic analysis. As the noise levels increase, the two states merge, and we find a single peak in the distribution of gene expressions for noise levels between 49 and 50 (the histogram on the right in Fig. 3A). This observation of state merging can be explained as follows. When the noise increases, the contribution of noise on gene expression exceeds the contribution of the regulatory interactions. Therefore, the circuit under high noise does not have the two distinct states anymore; instead, the only state left has similar expression of both genes. Since the two clusters are symmetric, both of the MIC-based and SA-based methods produce the same results.

**Fig. 3:**
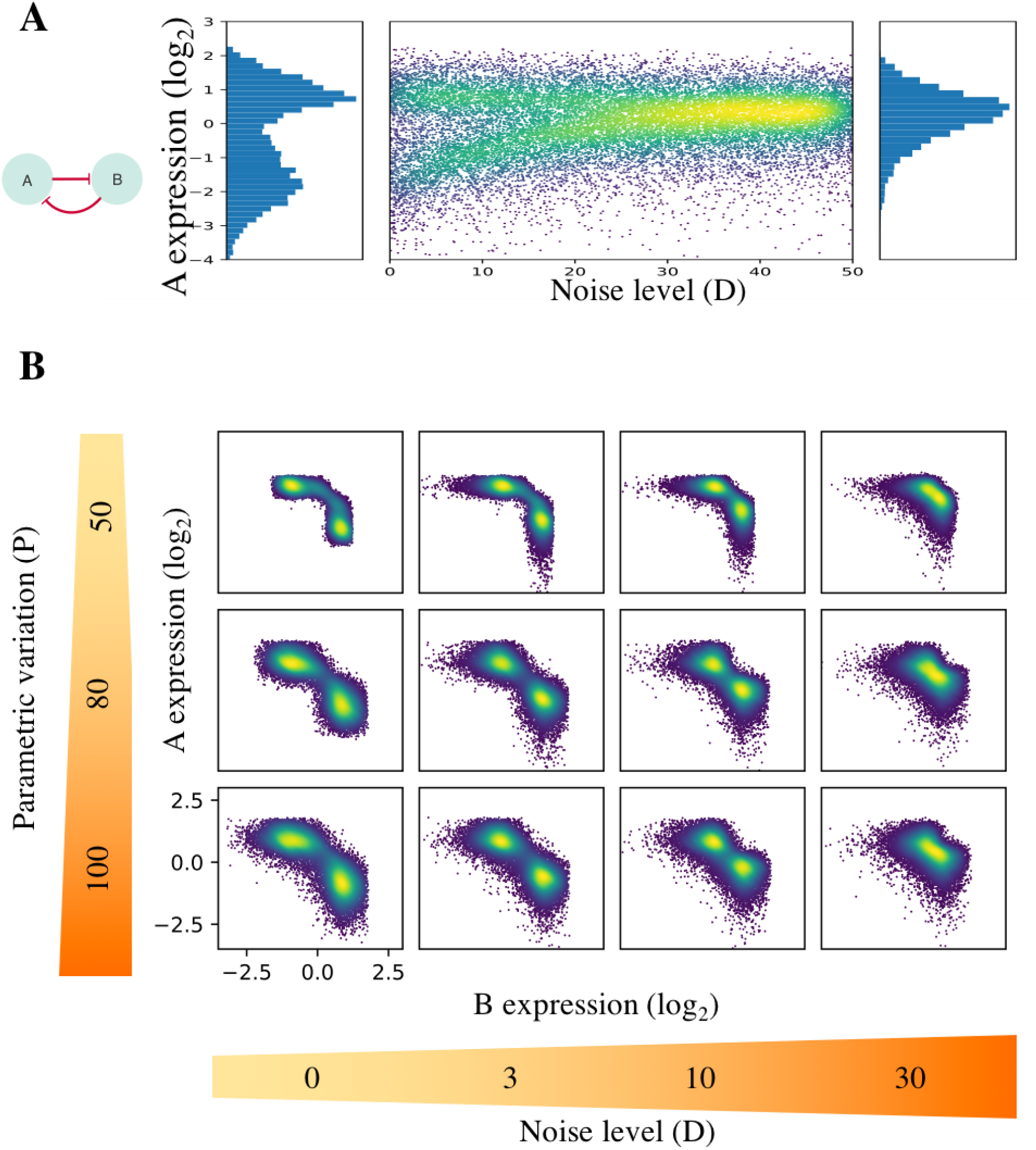
Application of sRACIPE to a toggle switch circuit. The stochastic analysis was performed using the MIC scheme. (A) The circuit diagram is illustrated in the leftmost panel. The middle panel shows the steady-state expression levels of gene A at different noise levels. The histogram on left shows the two distinct states for low noise (D<1), and the histogram on the right shows a single state for high noise (49<D<50). (B) Heatmaps of the normalized gene expression levels of the two genes for different parametric variations and noise levels. The parametric variations increase from top to bottom and the noise levels increase from left to right. Two distinct clusters observed at low noise merge into a single new cluster at high noise. Parametric variations increase the spread of the clusters but do not affect their relative positions.

Here, we treated the noise level as a control parameter and evaluated the response of gene expression. In a sense, this analysis can be considered as a *global bifurcation analysis*. Unlike traditional bifurcation diagram, where one alters a single parameter and keeps the other parameters constant, this global bifurcation analysis considers variations from the other parameters as well. Thus, this method has the potential to provide global pictures of systems under the control of a parameter, which in this case is the noise level. Similar idea can be applied to any other parameter (Fig. S3).

### Differential roles of noise level and parameter variation

We further explored the behavior of the toggle switch GRC by changing both the noise levels and parametric variations. Here, the parametric variation (P) is defined as the spread of the parameter ranges relative to the parameter range used in the original RACIPE^20^ while keeping the median constant. P is measured in percentages such that the ranges are same if P is set to 100, and a larger P implies a wider spread of parameter values. For any given value of P, if the range of a parameter is set to be (x_min_, x_max_) by default in RACIPE, the new range (y_min_, y_max_) can be obtained as

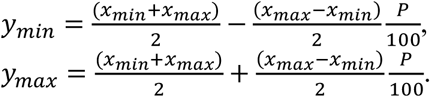

Using both the noise levels and parametric variations as two control parameters, we can plot a global 2D bifurcation diagram, as shown in Fig. 3B. We observe that while an increase in the parametric variations increases the spread around the two states, an increase in noise levels brings the states closer, and eventually, for large noise levels, the two states merge. This new state is different from the two states obtained from the deterministic analysis (when noise is zero) and corresponds to the previously unstable state in which both genes are expressed. These results are consistent with previous studies in that gene expression noise can create new states of a GRC ^5,10,12,23,34–38^. We demonstrated this point by sampling a large space of parameters and systematically testing the circuit behaviors. Moreover, our results indicate differential roles of the parametric variation and expression noise in influencing circuits’ behavior.

### Application of sRACIPE to complex GRCs

Next, we studied some complex circuits, *i*.*e*., a toggle switch with one self-activation link, a toggle switch in which both genes are self-activating, and a circuit with five coupled toggle switches (Fig 4). Similar to the earlier toggle switch example, the number of states as well as the gene expression pattern of these states changes with the increase in noise levels. These circuits have more than two states (i.e. clusters) and different states merge at different noise levels, suggesting these states have different levels of stability. For example, in the toggle switch with self-activations on both genes, the third intermediate cluster merges before the merging of two larger clusters. We used both of the MIC and SA methods to evaluate the basins of attraction and the stability of the states. Similar to the double well potential cases discussed earlier, we observe that the number of models in the different states at high noise is similar for both MIC and SA (again indicating that both methods estimate the stability), and more stable states have larger number of models. At low noise, the difference between the two methods can be observed prominently for the toggle switch with one self-activation, indicating different basins of attraction and stability of the two states. The difference is less evident in the symmetric cases where the two dominant states are not affected much, but the intermediate state has lesser number of models at low noise using the SA method. In short, sRACIPE can provide a global view of the dynamics of GRCs and allow the estimation of the basin of attraction by MIC and the stability by SA.

**Fig. 4:**
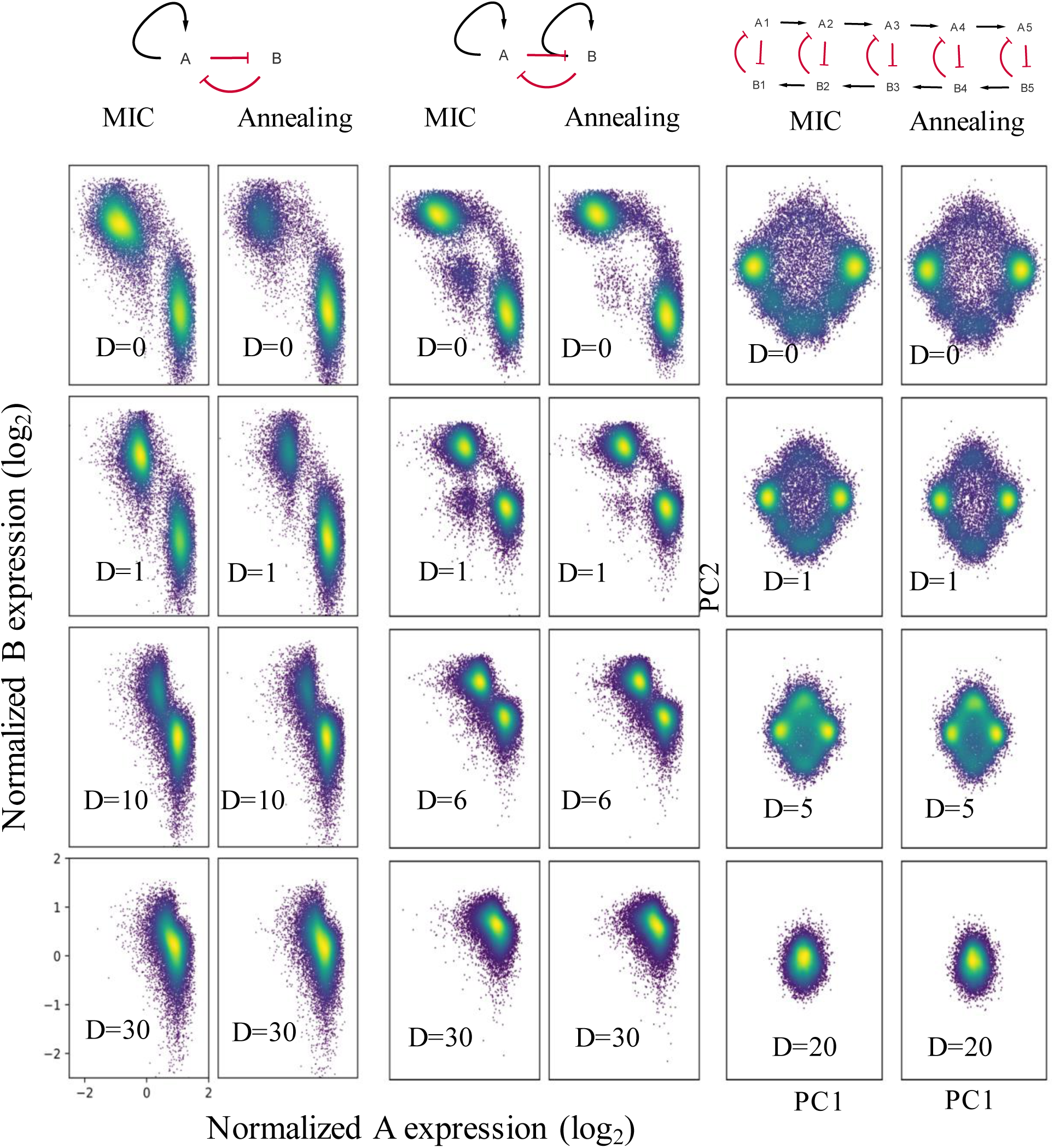
Normalized gene expressions for several toggle-switch-like-circuits using the MIC and SA methods. When noise is low, the MIC scheme provides an estimate of the basins of attraction of the states, whereas SA provides the most stable state. At high noise, the two methods yield similar results. The results are presented for (A) a toggle switch in which one gene is self-activating, (B) a toggle switch in which both genes are self-activating and (C) a circuit with five coupled toggle switches. Principal component analysis of the gene expression patterns was used for dimensionality reduction and the first two components are shown here. In all the cases, increase in noise levels brings the clusters together and eventually merges them into a final state. Some clusters merge first, suggesting that they are less stable than the others.

### Quantification of GRC’s robustness

We also observed that noise improves system’s response time, or so as to say, the time that the circuits take to reach the steady state probability distribution decreases with the increase in the noise levels (Fig. 5 for the results of the toggle switch GRC). Here, we compared the probability distributions at multiple time points to the probability distributions at the end of the simulations by calculating the Bhattacharyya Distance (BD, details in SI Methods) between them. Saturation in the BD values implies that the system has relaxed and converged. At higher noise levels, there is more variability in the steady state distributions, so the saturated BD values are larger for higher noise levels. But the system reaches this saturated BD value at a shorter simulation time. Further, we found that self-activating switches have larger BD than switches without self activations (Fig 5B), indicating that circuits with self-activating loops are less robust against noise. To quantify the robustness of GRCs against noise, we define the *noise robustness* (R_D_) index of a GRC as the rate of the increase of BD with the increase in noise level in the low noise limit:

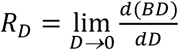

**Fig. 5:**
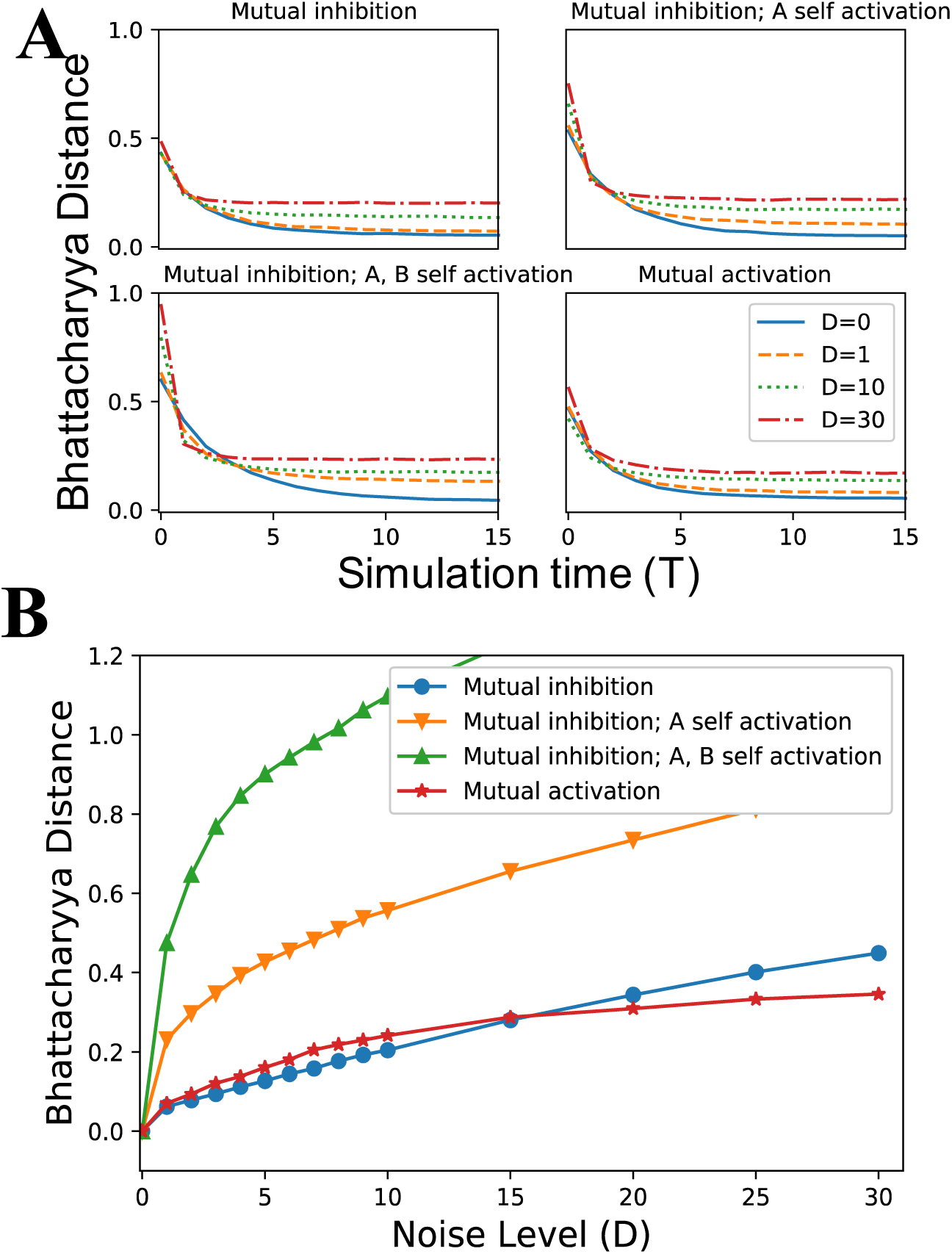
Response time and noise robustness of gene regulatory circuits. (A) Tests were performed on a toggle switch circuit. The Bhattacharyya distance (BD) was calculated between the probability distribution of the gene expression sampled at the end of the simulations (simulation time, T=50) for 10^6^ models at zero noise (D=0) and the probability distributions at different time points during the simulations for different noise levels. The BD value for D=0 approaches zero after T=15, indicating that the models have converged to steady state solutions. Simulations with larger noise converge faster, though BD is larger. (B) The response of BD with respect to the noise levels for different toggle-switch-like circuits. The results indicate that the self-activation links decrease the noise robustness.

The R_D_ values for the toggle switch, the toggle switch with one self-activation, the toggle switch with two self-activations and two mutual activating genes were found to be 0.06, 0.23, 0.48 and 0.07, respectively. We expect R_D_ to be valuable to quantify the robustness and stability of a gene circuit.

### Application to a GRC governing EMT

Lastly, we applied sRACIPE to an EMT gene regulatory circuit, which we previously constructed from former studies and an extensive literature search^20^. Evidences^39–42^ suggest that the EMT circuit controls the decision making of the cell transition from the epithelial to mesenchymal states during embryonic development, wound healing and cancer metastasis^40,41,43^. Hybrid epithelial/mesenchymal (E/M) states^25,41^ with mixed characteristics of collective cell migration, as in morphogenesis and wound healing, have been found in experiments^40^ as well as several computational modeling studies^25,41^, including our previous RACIPE analysis^20^.

The EMT GRC consists of 9 microRNAs (miRs) and 13 transcriptional factors (TFs), as shown in Fig. 6. In the deterministic case, clustering of the steady states of the randomized models yields two big clusters and other small clusters^20^ (also in Fig.6C, the first row). In the first big cluster, CDH1 and miRNAs are highly expressed, whereas some other TFs like ZEB1, ZEB2, CDH2, SNAI1, and SNAI2 are not expressed. This state can be mapped to the epithelial state (E). Similarly, in the other big cluster, miRNAs are not expressed, and TFs are highly expressed. Thus, it can be mapped to the mesenchymal state (M). The other clusters are hybrid states in which these key TFs and miRs have intermediate expressions. Here, we study how different levels of noise in the system affect the gene expression patterns of the models. The expression levels of different genes across all models vary over a large range; therefore, we scaled the noise level of each gene by the median of the expression of that gene over all models in the deterministic case. We observed relative shifts in the E and M states and more models in the hybrid E/M states as the noise levels increase. Eventually, the clusters merge together when noise levels became very high. Comparing the heatmaps for D=0 and D=10^−5^, we notice an increase in the number of models in the E state and the hybrid states. Notably, the CDH1 levels have increased significantly even in the models corresponding to the M state. On further increase of the noise levels, the expression levels of miR-200b increase (D=10^−3^), and TGF-beta decrease in the M (D=10^−1^) state. Therefore, CDH1, miR-200b and TGF-beta are susceptive to noise perturbation, suggesting they may play crucial role in EMTs. Indeed, this is in line with the earlier findings that these genes are crucial in distinguishing the EMT phenotypes^20,39^. We have explored possible mechanisms to stabilize the hybrid EMT phenotype in our previous studies^41,44^. Here we present an additional mechanism in which the hybrid EMT phenotype can be stabilized due to the increase of gene expression noise. It would be interesting to validate this hypothesis experimentally in the further. In addition, our SA simulations suggest that the E state is more stable than the M state (Fig.6B).

**Fig. 6:**
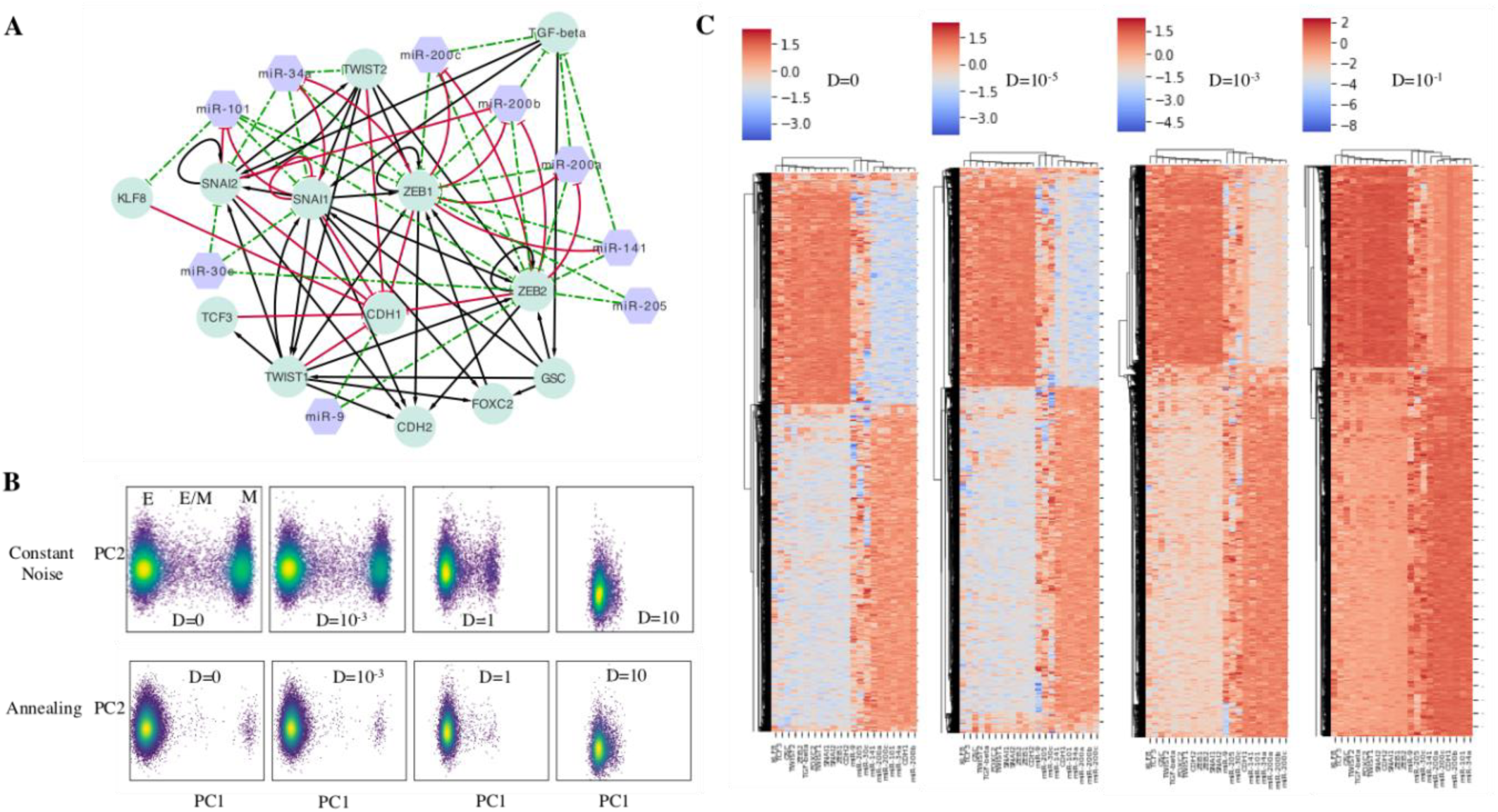
Application of sRACIPE to an EMT core circuit. (A) An EMT core circuit consisting of 9 microRNAs (miRs) and 13 transcription factors (TFs). The activating links from a TF are shown in black, the inhibitory links from a TF are shown in red and the regulatory links (inhibitory) starting from miRs are shown in green. (B) The first and second principal components from the gene expression patterns at different noise levels for both of the MIC and SA schemes of sRACIPE. Here, the noise level in each gene is scaled by its median expression from the deterministic analysis. The steady states corresponding to the E and M states form two distinct clusters in the PC1/PC2 projection, with the hybrid states occupying the space between them. With the increase in noise levels, both of the E and M states move toward each other and finally reach the hybrid states. The shifting of the M state is larger than that of the E state. From the simulations of the SA scheme, we find that the M state is much less stable than the E state. (C) Hierarchical clustering heatmaps using the MIC scheme. As the noise levels increase, the models in the M state have increased CDH1 expression, followed by increased miR200b expression.

## Discussion

In this work, we have developed a method, named sRACIPE, to integrate stochastic analysis with the random circuit perturbation (RACIPE) method. It allows us to study the effect of both gene expression noise and parametric variations on any gene regulatory circuit (GRC) using only its topology. To facilitate sampling, we proposed two ensemble-based schemes for stochastic analysis. The two methods, constant noise simulations with multiple initial conditions (MIC) and simulated annealing (SA), complement each other to provide a holistic picture, where MIC estimates the basin of attraction and SA estimates the stability. Our tests show that expression noise and parametric variation have qualitatively different effects on the states of GRC. Parametric variation slightly broadens the spread of the states while high expression noise causes some states to merge together. We have found that GRCs with different topology have different response times and sensitivity to noise. We have implemented sRACIPE as an R package, and it will be freely available for academic use.

By sampling only one initial condition for each model, sRACIPE can easily generate as many as 10^6^ models. One major challenge is how to fully utilize such a large amount of gene expression and parameter data to analyze the robust features of a GRC. These data analysis methods can be potentially used to quantify the robustness of a GRC and evaluate how this can be associated with evolutionary fitness^45^, estimate the Waddington’s epigenetic landscape^46^, and predict state transitions^33^. A better understanding of stochastic behavior can be exploited to induce desired cell states and control noise-induced switching between different states^47^.

Both gene expression noise and parametric variations are common in biological systems^1,4,6,7,13,22,48^. On the one hand, the Gillespie algorithm (Fig S2) has been used to model the stochastic dynamics of gene expression caused by low copy number and slow switches between gene states^16^. On the other hand, cells of different size and microenvironment can be modeled by the same rate equations but different kinetic parameters^49^. Our method allows the analysis of both factors, therefore being an invaluable tool to study the nature of variations in a cell population, especially with the advent of single cell techniques.

We have found that GRCs with different circuit topology may allow similar states but differ in their sensitivity to noise, consistent with several theoretical and experimental studies^2,50^. Biological circuits are usually robust against small noise; sometimes, they could even use noise for their functions^1^. For example, noise can create new states or destabilize existing ones^5,10,12,23,34–38^.

In summary, we have presented an improved randomization-based method for gene circuit modeling by incorporating stochastic analysis. This method is relevant to the study of multi-stable biological processes. Our methods provide a quantitative characterization of the robustness of biological networks in the presence of both intrinsic and extrinsic noises. Further investigation of biological data with the modeling approach is expected to provide better insight into the role of noise in network dynamics.

### A public R package of sRACIPE

We provide an R package of sRACIPE, which is freely available for academic use. The core program is written in C++ to speed up the simulations, and the interface of R facilitates the processing of user inputs. We also provide R scripts to perform the basic statistical analyses that we have mentioned in the paper. We hope sRACIPE will be a valuable systems biology tool to analyze gene regulatory circuits.

## Acknowledgement

The study is supported by a startup fund from The Jackson Laboratory. Mingyang Lu is partially supported by the National Cancer Institute of the National Institutes of Health under Award Number P30CA034196.

## Supplementary information

S1: Supplementary methods and figures

S2: Simulated annealing in a double well potential (movie)

S3: EMT topology file

